## Supplementary data for "Scube2 primes Dispatched and ADAM10-mediated Shh release by recruiting HDL acceptors to the plasma membrane"

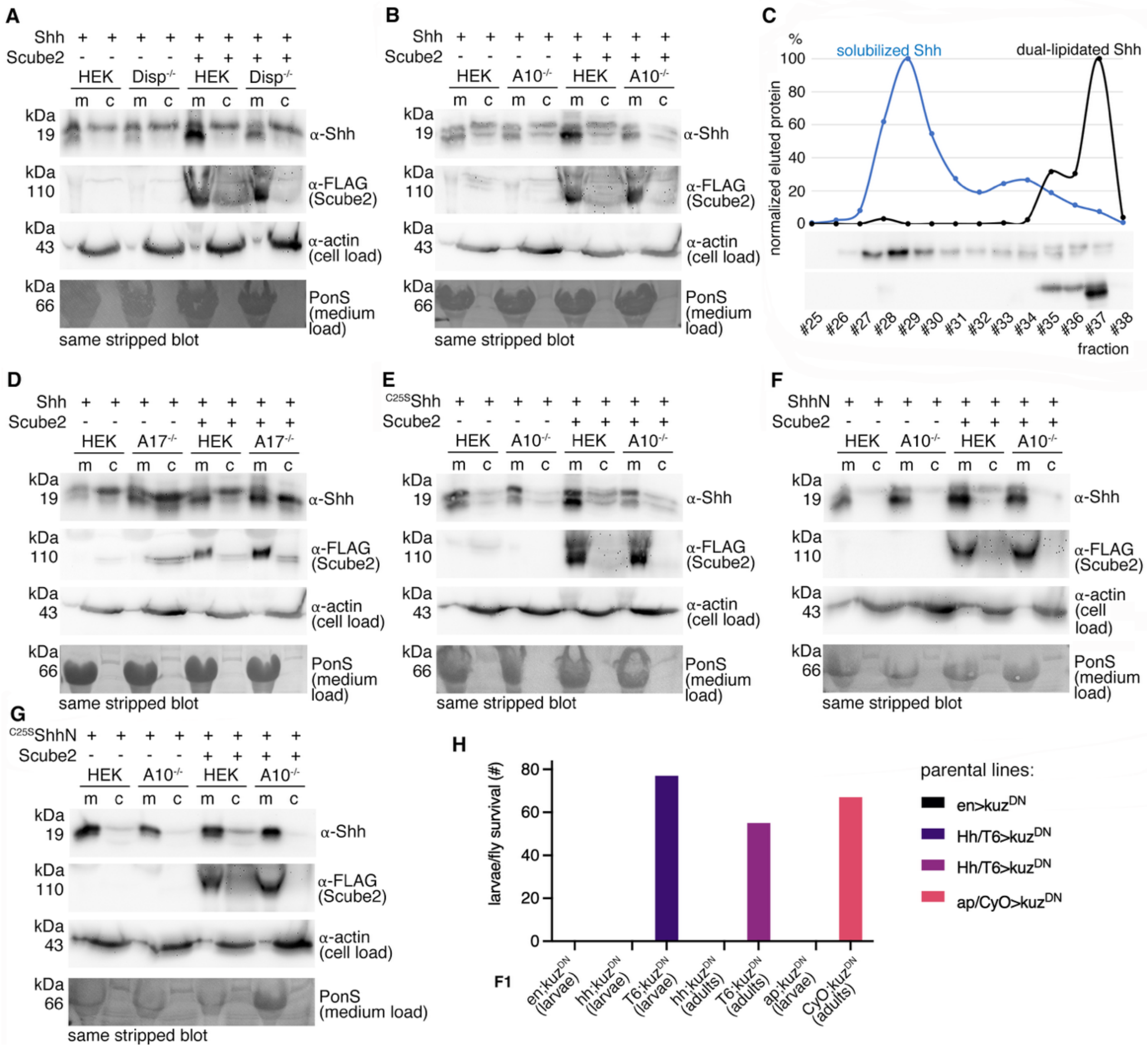

**Supplementary Figure 1, related to Figures 1-2. Loading controls, activity controls and standards. A, B)** In this and subsequent figures, α-Flag antibodies detected Flag-tagged Scube2, actin served as a loading control for cellular lysates, and PonceauS (PonS) served as a loading control for serum BSA traces in media in this and subsequent figures. Note that the amounts of soluble and cellular Shh in this and subsequent figures are not inversely correlated. This is because the medium lanes represent all TCA-precipitated proteins in the medium, while the cells were lysed directly in SDS buffer and only a small fraction (approximately 5%) was applied to the gel. As a consequence, a 100% increase in Shh solubilization will correlate to only 5% reduction in the amount of cell-surface-associated Shh, and *vice versa*. **C)** RP-HPLC of solubilized Shh from HEK293 cells (blue) and commercial dual-lipidated R&D 8908-SH Shh, representing the dual-lipidated precursor protein at the cellular surface of producing cells (black). The elution profiles of dual-lipidated cellular Shh and the solubilized protein do not overlap, demonstrating Shh delipidation during release. For more information on Shh delipidation during release, also see [1]. **D)** A17 does not contribute to Shh solubilization in the presence of Scube2. Note increased rather than decreased Shh solubilization from $A17^{-/-}$ cells, which may be explained by compensatory upregulation of another protease [2] to cleave Shh, possibly A10. One representative of 5 experiments with similar results is shown. **E-G)** Actin, PonceauS, and Flag-tagged Scube2 loading controls for the immunoblots shown in Figure 2. **H)** $Kuz^{DN}$ expression under the control of Engrailed (en), Hedgehog (hh), and Apterous (ap) leads to early embryonic (en, ap) or L1 larval (hh) lethality. This prevented the analysis of Kuz-regulated Hh release in *Drosophila* wing discs.

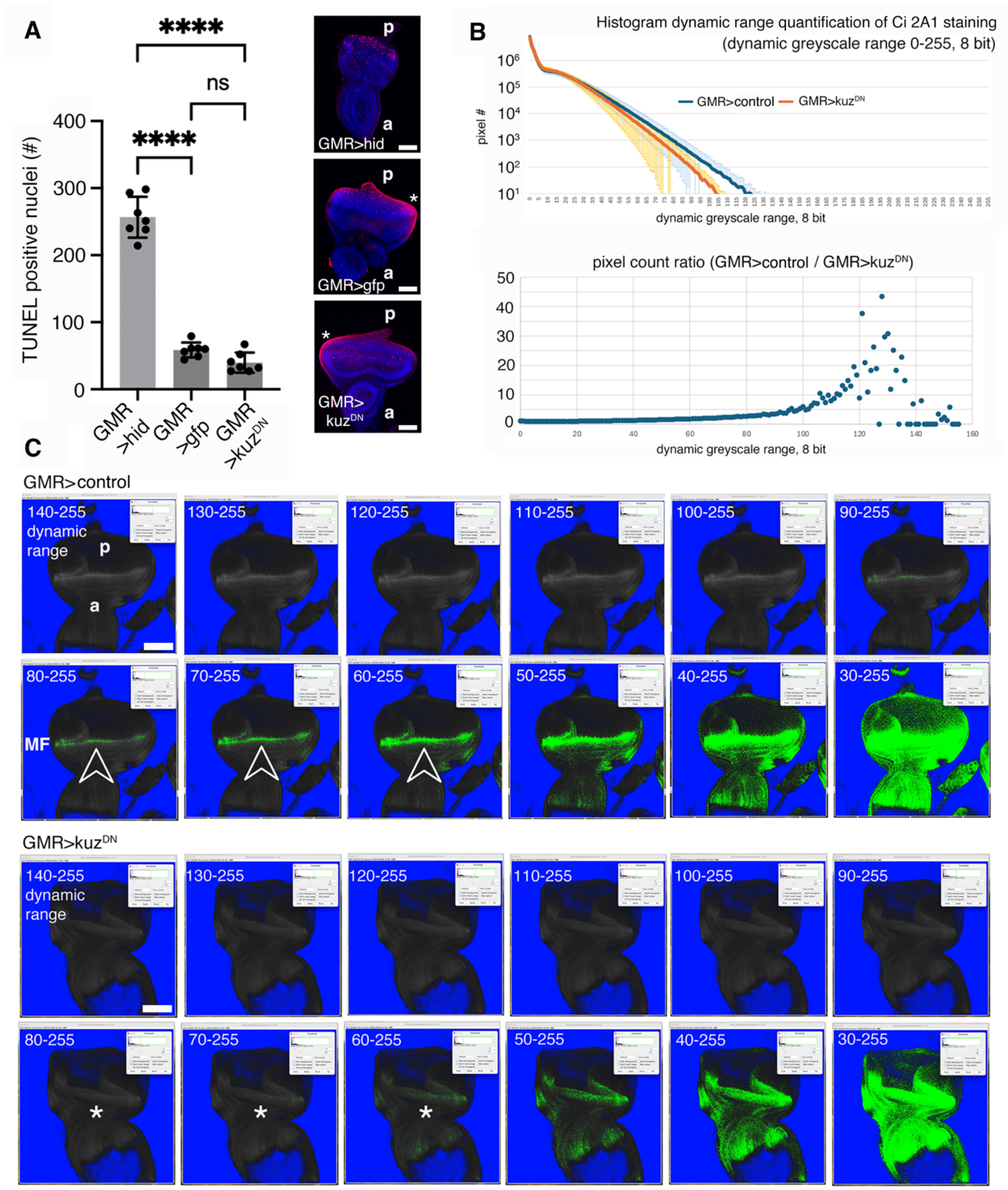


**Supplementary Figure 2, related to Figure 3. Reduced Ci expression, rather than increased apoptosis, underlies the small eye phenotype in flies expressing kuz$^{DN}$ under GMR control. A)** DNA nick end labeling (TUNEL) of L3 eye discs. Apoptosis in discs expressing kuz$^{DN}$ under GMR-Gal4 control was not increased compared to mCD8-GFP expressing control discs (GMR>gfp). In contrast, apoptosis in GMR>hid positive control eye discs was strongly increased, confirming proper assay conditions. ****: p<0.0001; n.s.: 0.23, n = 7. One representative disc of each genotype is shown, asterisks indicate unspecific staining at the tissue border. a: anterior, p: posterior ends of the discs. Scale bar: 50µm. **B)** Histogram dynamic range quantification of ci 2A1 staining (top) and pixel count ratio applied to the same dataset. Anti-Ci-stained L3 eye discs from larvae expressing FLP (n=7 discs) or Kuz$^{DN}$ (n=8 discs) under GMR control demonstrate decreased Ci expression dynamics as a consequence of impaired Hh solubilization in $Kuz^{DN}$-expressing discs. The pixel count ratio of the same dataset emphasizes that the intensity of the highest signals is diminished in the $Kuz^{DN}$-expressing discs, while the base-level intensity remains unaffected. **C)** Dynamic range variation reveals a peak Ci staining (shown in green) at the morphogenetic furrow (MF) of GMR>FLP control eye discs (arrowheads) close to the Hh-releasing nascent photoreceptor cells. Using the same settings, the series of GMR>$kuz^{DN}$ discs shows an evident lack of such a peaking Ci staining (asterisks). The dynamic range of Ci expression in the antennal part of the discs is unaffected by $Kuz^{DN}$, which serves as an internal calibration control (bottom right) for Ci intensities at the MF. Scale bars: 50µm. These results are consistent with the reduced release of endogenously expressed Hh from nascent photoreceptors in discs with diminished endogenous Kuz function. This impaired release, in turn, reduces Hh signaling and downstream Ci expression in adjacent receiving cells at the morphogenetic furrow. The numbers indicate the dynamic range set, which is a subrange of the total 8-bit, 0–255 greyscale range, to clearly demonstrate the quantitative Ci expression differences at the furrows of the two genotypes.

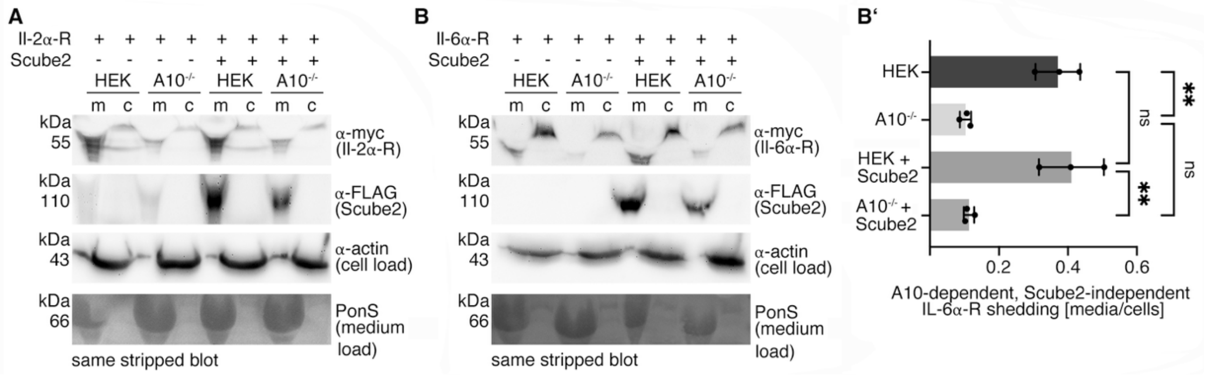


**Supplementary Figure 3, related to Figure 4. Scube2 is not a general A10 activator. A)** HEK control or $A10^{-/-}$ cells were seeded on day 0 and transfected with IL2α-R – a known A10 substrate – together with Scube2 or empty control cDNA3.1. Cells were grown in DMEM containing 10% FCS for 36 h, serum containing media was aspirated and serum-free DMEM was added for 6 h (representing the serum-depleted conditions used in previous experiments). Media were collected and centrifuged at 300g for 10 min to remove debris, 10% trichloroacetic acid was added, proteins were precipitated on ice for 30 min, followed by centrifugation at 13,000g for 20 min. Cells were lysed directly on the plate for subsequent SDS-PAGE analysis. In contrast to what we observed for Shh, IL2α-R release was dependent on A10 (confirming the knockout cell type used in our experiments) but not on Scube2. The figure includes loading controls for actin, total soluble proteins and Scube2 for the immunoblot shown in Figure 4A. **B)** Transfection of HEK control cells or $A10^{-/-}$ cells with IL6α-R, another established A10 substrate, together with Scube2 or empty control cDNA3.1. Actin, PonceauS and Flag-tagged Scube2 loading controls are also shown. Like IL2α-R, IL6α-R release was dependent on A10 but not on Scube2. **B')** Quantification of B. **: $p<0.002$, n.s.: $p>0.05$.

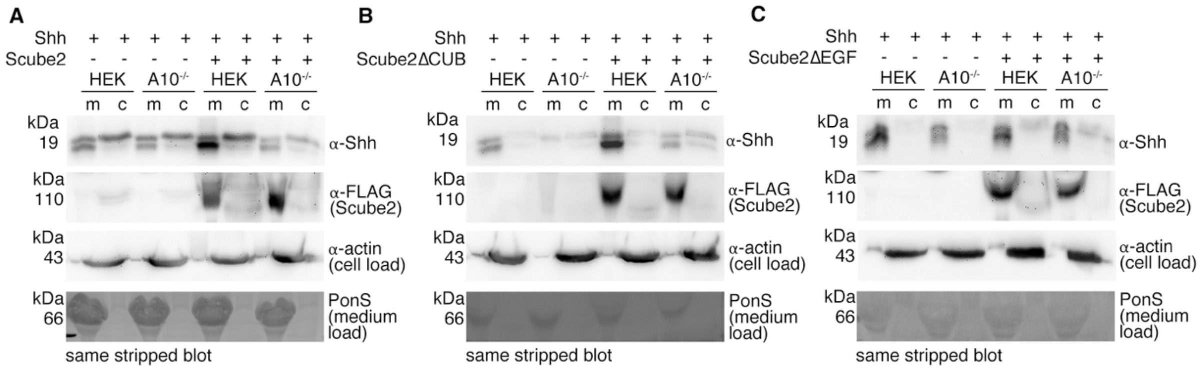


**Supplementary Figure 4, related to Figure 5. Loading controls. A-C)** Actin, PonceauS, and Flag-tagged Scube2 loading controls for the immunoblots shown in Figure 5.

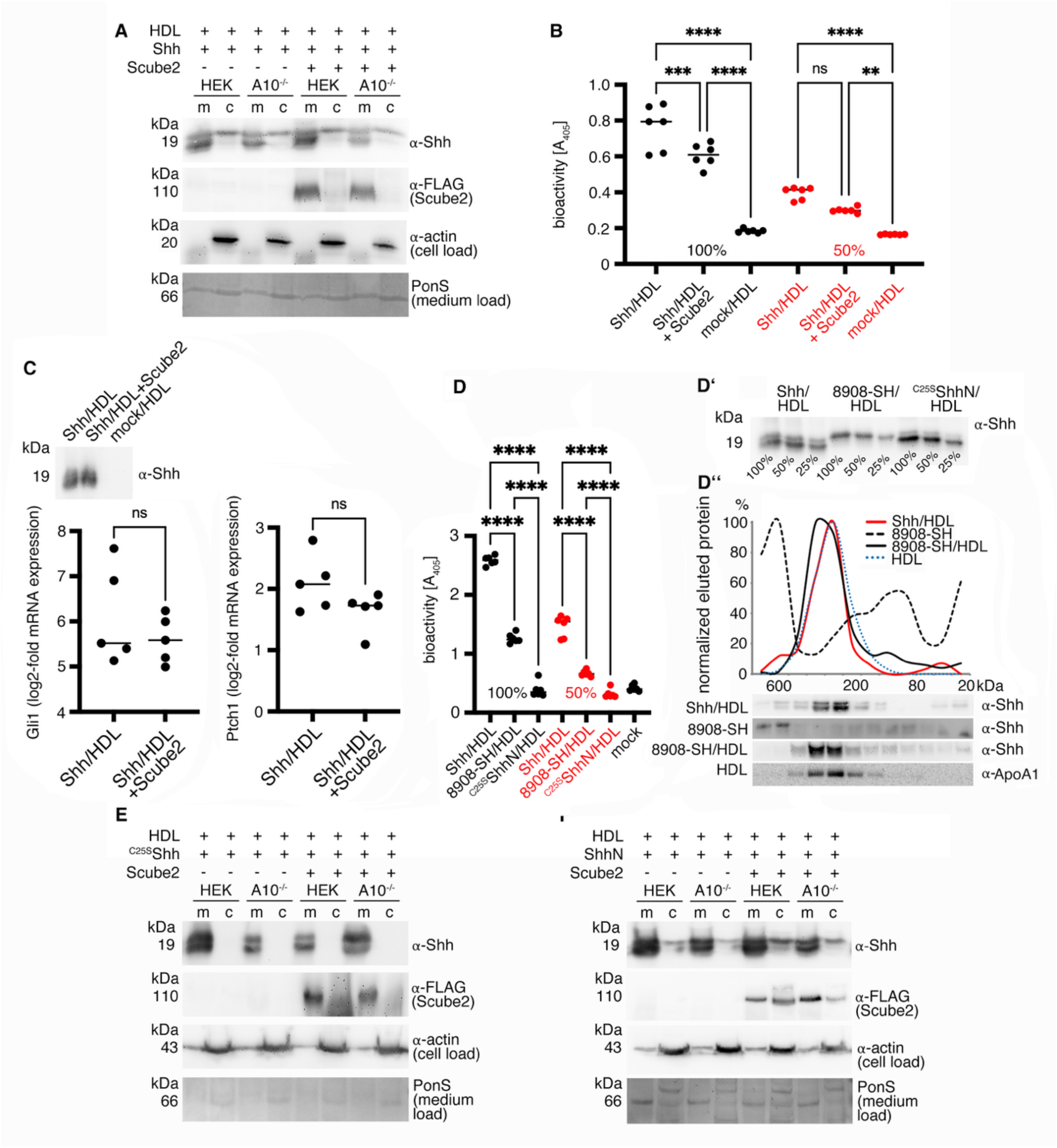

**Supplementary Figure 5. Loading controls and bioactivity assays. A)** Shown are actin, PonceauS, and Flag-tagged Scube2 loading controls for the blots shown in Figure 6B. One out of four independent experiments with similar results is shown. **B)** N-processed (depalmitoylated) Shh released on HDL is bioactive. Shh with or without co-transfected Scube2 was solubilized from washed Disp- and A10-expressing cells in serum-free media containing 40µg/ml HDL. The harvested medium was adjusted to a final concentration of 10% FCS and added to C3H10T1/2 reporter cells at two different concentrations to assess the possibility of oversaturation of the system with bioactive Shh. Both HDL-associated Shh samples induced C3H10T1/2 reporter cell differentiation in a concentration-dependent manner as determined by Shh-induced alkaline phosphatase (Alp) activity measured at 405 nm. **C)** N-processed depalmitoylated, HDL-associated soluble Shh induced similar increases in the Shh target genes ptch1 and gli1 in stimulated C3H10T1/2 osteoblast progenitor cells, regardless of the presence or absence of Scube2. Gene expression changes are expressed relative to levels induced by HDL-containing media from mock-transfected cells. Both results demonstrate that Scube2 is not required as a carrier for Shh to the Ptch receptor if linked to HDL and that the Shh palmitate is not required for signaling at receiving cells under this condition. Similar amounts of solubilized Shh, as determined by immunoblotting, were used in both assays (inset). **D)** N-processed, HDL-associated soluble Shh induced C3H10T1/2 reporter cell differentiation in a concentration-dependent manner, as determined by Shh-induced Alp activity measured at 405 nm. Dual-lipidated Shh (R&D 8908-SH), which represents the cell-surface-associated form upstream of Disp-, A10-, and Scube2-mediated release, was mixed with the same amount of HDL used for Shh release from cells and added to C3H10T1/2 reporter cells. The significantly reduced bioactivity of the dual-lipidated 8908-SH variant supports the previously published idea that the unprocessed Shh N-terminal peptide sterically inhibits Ptch receptor binding [3] and signaling, and that proteolytic removal of the N-peptide renders Shh accessible to Ptch [4]. Unlipidated $^{C25S}$ShhN known to be devoid of signaling activity and HDL only (mock) served as a negative controls. ****: $p<0.0001$, $n=6$. **D')** Western blot demonstrating similar amounts of Shh in the conditioned media used for this experiment. **D'')** Size exclusion chromatography (gel filtration) analysis of the same material shows a wide molecular weight distribution of dual-lipidated, detergent-solubilized 8908-SH in the absence of HDL (black dashed line), but not in its presence (black solid line). This shows that the protein rapidly associated with the LPP, resulting in HDL-associated clusters that were very similar to those resulting from Shh release from Shh transfected cells in the presence of HDL (red line). However, their activities differ significantly, as shown in D. The blue dotted line shows the HDL signal from the same blot. **E, F)** Solubilization of non-palmitoylated $^{C25S}$Shh and non-cholesteroylated ShhN in serum-free media supplemented with 40µg/ml HDL. $^{C25S}$Shh and ShhN release is independent of A10 and Scube2, demonstrating that dual Shh lipidation is essential for physiologically relevant transfer to HDL.

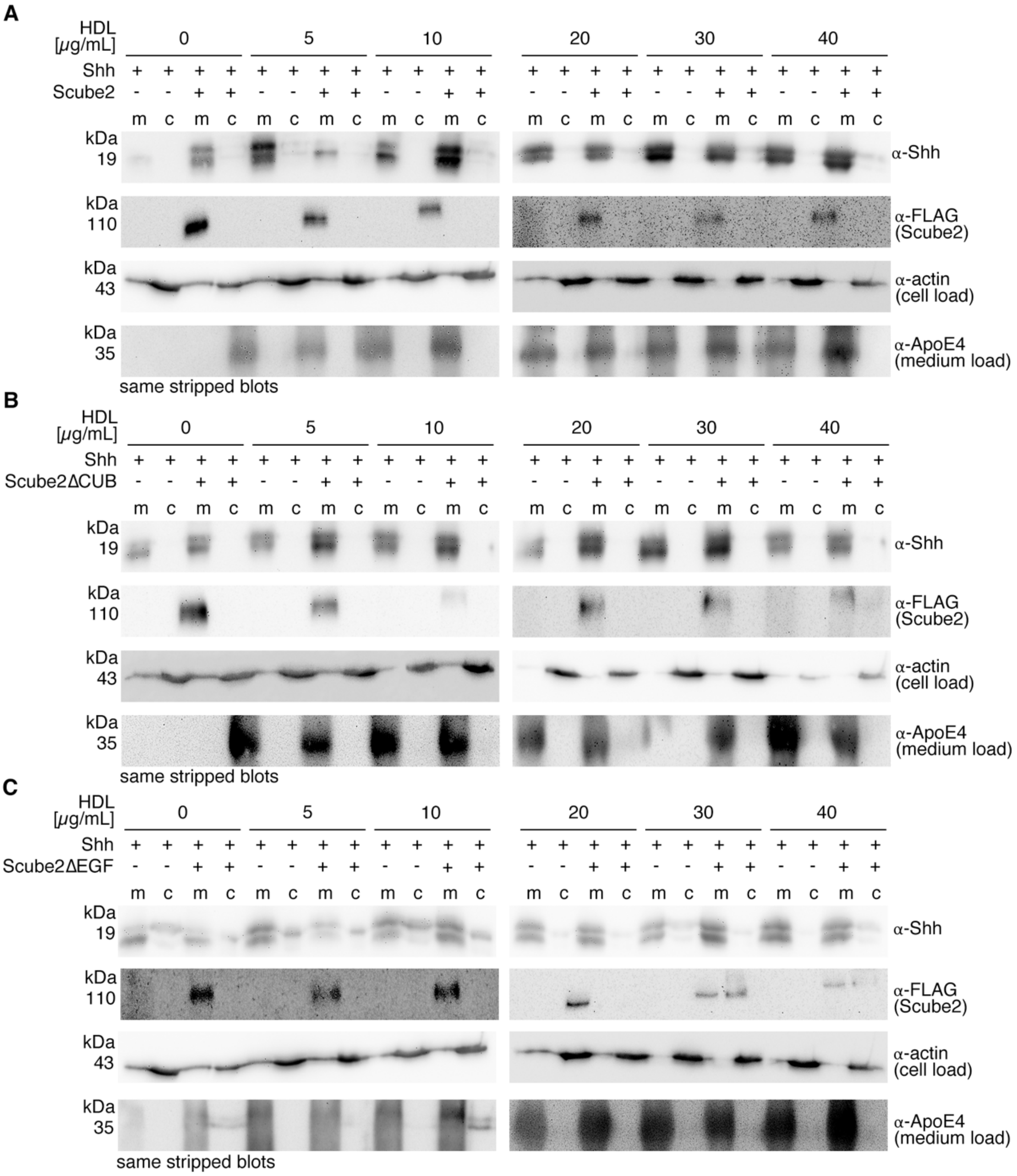


**Supplementary Figure 6. Loading controls related to Figure 6D, E.** Scube2 and Scube2ΔCUB, but not Scube2ΔEGF increase Shh solubilization when HDL concentrations are low. Actin, HDL (detected using antibodies directed against ApoE4) and Flag-tagged Scube2 loading controls are shown.

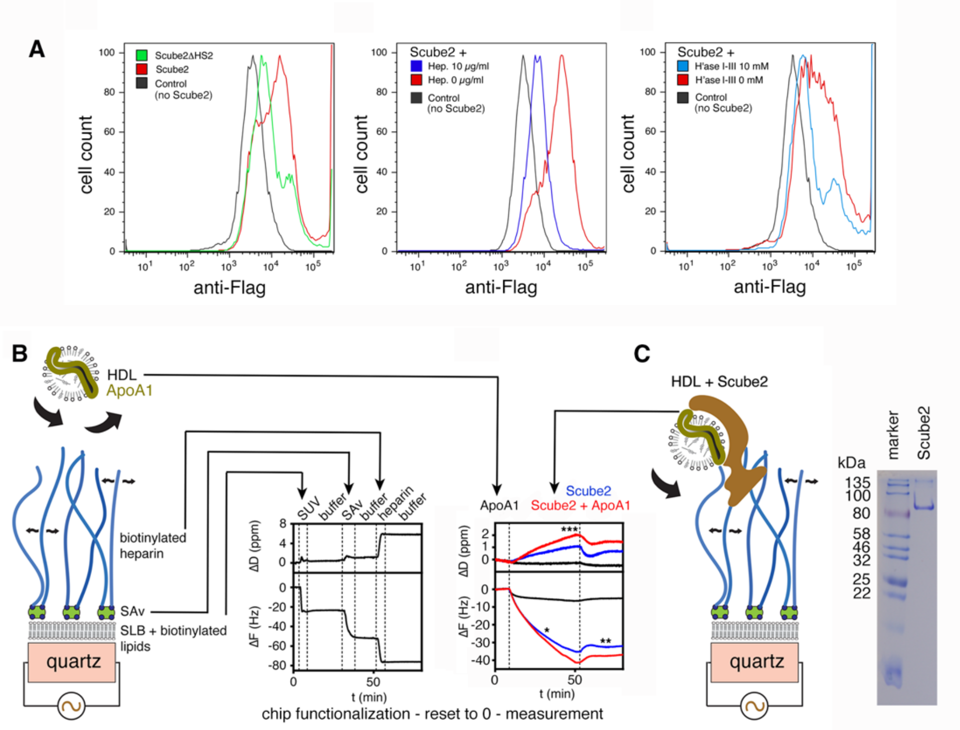


**Supplementary Figure 7: Scube2 binding to HS and heparin, as determined by Fluorescence Activated Cell Sorting (FACS) and Quartz Crystal Microbalance with Dissipation Monitoring (QCM-D). A)** FACS of Scube2-expressing Bosc23 cells reveals that Flag-tagged Scube2 associates with the cell surface of HEK293 cells (red line) when compared with control cells without Scube2 (grey line). Scube2 specifically binds the HS expressed at the HEK293 surface, because surface binding of a Scube2 variant lacking one main motif of HS-binding basic amino acids (ΔHS2, green line) [5] is strongly reduced. Soluble heparin (10 µg/ml) also specifically competed with Scube2 binding to the cell surface (center, blue line), and cell-surface HS degradation by 7.5 mU heparinases I, II and III (2.5 mU each) impairs Scube2 association with the cell surface as well (right). A total of 50,000 cells were analyzed under each condition. **B)** The core of the QCM technology is a gold-coated oscillating quartz crystal sensor disc with a resonance frequency related to the mass of the disc. Interaction surfaces containing biotinylated heparin (blue) linked via steptavidin (Sav, green) to fluid-supported lipid bilayers (SLBs, black) were generated as a proxy for HS at the surface of Shh producing cells. Like cell surface HS, SLB-linked heparin, which is highly negatively charged, can freely rotate and move laterally on the sensor surface (curved arrows). QCM-D allows real-time detection of nanoscale mass changes on the sensor surface (binding or unbinding of soluble molecules to the disc) by monitoring changes in resonance frequency ($\Delta F$). QCM-D measures an additional parameter, the change in energy dissipation $D$, which is particularly useful for studying the viscoelastic properties of the surface layer, which tend to increase when a soluble ligand is bound. Shown on the right are representative QCM-D data showing the observed frequency ($\Delta F$) and dissipation ($\Delta D$) shifts during the assembly of the SLB, the SAv monolayer, and the film of terminal heparin on the silica surface of the sensor. The formation of the heparin/HS model matrix on the QCM sensor was always followed in real time prior to the protein incubation assays shown in Figure 6F to validate the surface functionalization, and $D$ and $F$ were reset to zero prior to the actual measurement. **C)** After reset, ApoA1 alone or together with Scube2 were added to the buffer that washes the sensor. The start and duration of sample incubations are indicated by dashed vertical lines and a label at the top of the graph. At all other times, the surface was exposed to wash buffer. We observe that ApoA1 alone does not bind to the sensor surface, but Scube2 alone or with ApoA1 binds strongly, indicating its potential to recruit HDL to HS-rich cell surfaces (as shown in the cartoons). Outer right: Coomassie-stained purified Scube2 used in the QCM-D assays.

**Supplemental Table 1: Statistical source data**

| Figure | Cell type | Constructs | Values (mean) ± SD, n | p-values |
|---|---|---|---|---|
| 1B’ | Disp$^{-/-}$ / HEK | Shh + Scube2 | 0,3746±0,02056 vs 0,6254±0,0256, n=6 | <0,0001 |
| 1B’’ | HEK | Shh ± Scube2 | 0,2874±0,08772 vs 0,7143±0,08772, n=6 | <0,0001 |
| 1C’ | A10$^{-/-}$ / HEK | Shh + Scube2 | 0,1951±0,04876 vs 0,6348±0,1672, n=12 | <0,0001 |
| 1C’’ | HEK | Shh ± Scube2 | 0,2666±0,02818 vs 0,7334±0,02818, n=6 | <0,0001 |
| 2A’ | A10$^{-/-}$ / HEK | $^{C25S}$Shh + Scube2 | 0,3324±0,05179 vs 0,6676±0,0,5179, n=4 | <0,0001 |
| 2A’’ | HEK | $^{C25S}$Shh ± Scube2 | 0,3549±0,05269 vs 0,6451±0,05269, n=4 | 0,0002 |
| 2B’ | A10$^{-/-}$ / HEK | ShhN + Scube2 | 0,4609±0,06991 vs 0,5391±0,06991, n=6 | 0,0813 |
| 2B’’ | HEK | ShhN ± Scube2 | 0,4736±0,06644 vs 0,5264±0,06644, n=6 | 0,1984 |
| 2C’ | A10$^{-/-}$ / HEK | $^{C25S}$ShhN + Scube2 | 0,4820±0,06717 vs 0,5180±0,06717, n=8 | 0,3028 |
| 2C’’ | HEK | $^{C25S}$ShhN ± Scube2 | 0,4607±0,08287 vs 0,5393±0,08287, n=8 | 0,0787 |
| 3C | D.m. eye | GMR>gfp vs GMR>Kuz$^{DN}$ | 657,0±37,87 vs 235,1±21,81, n=10 | <0.0001 |
| 3C | D.m. eye | GMR>gfp vs hh$^{bar3}$/hh$^{bar3}$ | 657,0±37,87 vs 271,6±28,36, n=10 | <0.0001 |
| 3C | D.m. eye | GMR>gfp vs disp$^{LacW}$/disp$^{LacW}$ | 657,0±37,87 vs 304,5±57,69, n=10 | <0.0001 |
| 3C | D.m. eye | GMR>gfp vs GMR>$^{HA}$Hh; Hh$^{bar3}$/Hh$^{AC}$ | 657,0±37,87 vs 244,0±21,97, n=10 | <0.0001 |
| 3C | D.m. eye | GMR>gfp vs GMR>Hh; Hh$^{bar3}$/Hh$^{AC}$ | 657,0±37,87 vs 648,7±20,69, n=10 | 0,988 |
| 4A’ | A10$^{-/-}$ / HEK | IL-2Rα + Scube2 | 0,08572±0,03812, n=4 vs 0,4323±0,07046, n=6 | <0,0001 |
| 4A’ | HEK | IL-2Rα ± Scube2 | 0,44450,05869 vs 0,4323±0,07046 , n=6 | 0,9823 |
| 4A’ | A10$^{-/-}$ | IL-2Rα ± Scube2 | 0,09905±0,04566 vs 0,08572±0,03812, n=4 | 0,9873 |
| 4A’ | A10$^{-/-}$ / HEK | IL-2Rα | 0,09905±0,04566, n=4 / 0,4445±0,05869, n=6 | <0,0001 |
| 5B’ | HEK | Shh ± Scube2 | 0,2874±0,08772 / 0,7143±0,08772, n=6 | <0,0001 |
| 5C’ | HEK | Shh ± Scube2ΔCUB | 0,3934±0,07830 / 0,6066±0,07830, n=5 | 0,0026 |
| 5D’ | HEK | Shh ± Scube2ΔEGF | 0,4731±0,06399 / 0,5269±0,06399, n=6 | 0,1767 |
| 6B’ | A10$^{-/-}$ / HEK | Shh + Scube2 | 0,4195±0,04394 / 0,5805±0,04394, n=5 | 0,0004 |
| 6B’’ | HEK | Shh ± Scube2 | 0,4974±0,01995 / 0,5026±0,01995, n=5 | 0,6867 |

| | | | | |
|---|---|---|---|---|
| 6D | HEK | Shh ± Scube2 | 0,023±0,003 vs 0,133±0,025 (0 µg),<br>0,049±0,008 vs 0,089±0,011 (5 µg),<br>0,039±0,008 vs 0,08±0,011 (10 µg)<br>0,072±0,022 vs 0,083±0,027 (20 µg)<br>0,08±0,035 vs 0,085±0,032 (30 µg),<br>0,128±0,012 vs 0,139±0,016 (40 µg), n=4 | <0,0001<br>0,044<br>0,0425<br>0,9701<br>0,9993<br>0,9687 |
| 6E | HEK | Shh ± Scube2ΔCUB | 0,017±0,006 vs 0,113±0,052 (0 µg),<br>0,032±0,004 vs 0,094±0,005 (5 µg),<br>0,037±0,002 vs 0,094±0,03 (10 µg)<br>0,09±0,23 vs 0,133±0,028 (20 µg)<br>0,099±0,008 vs 0,094±0,005 (30 µg),<br>0,133±0,011 vs 0,104±0,008 (40 µg), n=4 | <0,0001<br>0,0009<br>0,0025<br>0,0319<br>0,9997<br>0,3098 |
| S2A | D.m. wing disk | GMR>hid (induced apoptosis) vs GMR>gfp (negative control) | 256,7±30,41 vs 58,57±11,12, n=7 | <0,0001 |
| S2A | D.m. wing disk | GMR>hid vs GMR>kuz$^{DN}$ | 256,7±30,41 vs 39,86±14,86 | <0,0001 |
| S2A | D.m. wing disk | GMR>gfp vs GMR>kuz$^{DN}$ | 58,57±11,12 vs 39,86±14,86 | 0,2315 |
| S3B’ | HEK | IL-6R ± Scube2 | 0,3716±0,06464 / 0,4110±0,09434, n=3 | 0,8399 |
| S3B’ | A10$^{-/-}$ / HEK | IL-6R + Scube2 | 0,1138±0,01489 / 0,4110±0,09434, n=3 | 0,0011 |
| S3B’ | A10$^{-/-}$ | IL-6R ± Scube2 | 0,1036±0,01617 / 0,1138±0,01489, n=3 | 0,9963 |
| S3B’ | A10$^{-/-}$ / HEK | IL-6R | 0,1036±0,01617 / 0,3716±0,0646, n=3 | 0,0022 |
| S5B | C3H10T1/2 | mock HDL vs Shh HDL (100%) | 0,2528±0,016 au vs 1,36±0,24 au, n=6 | <0,0001 |
| S5B | C3H10T1/2 | mock HDL vs Shh HDL + Scube2 (100%) | 0,2528±0,016 au vs 1,08±0,10 au, n=6 | <0,0001 |
| S5B | C3H10T1/2 | Shh HDL vs Shh HDL + Scube2 (100%) | 1,36±0,24 au ± 1,08±0,10 au, n=6 | 0,0005 |
| S5B | C3H10T1/2 | mock HDL vs Shh HDL (50%) | 0,218±0,003 au vs 0,654±0,06 au, n=6 | <0,0001 |
| S5B | C3H10T1/2 | mock HDL vs Shh HDL + Scube2 (50%) | 0,218±0,003 au vs 0,498±0,024 au, n=6 | <0,0001 |
| S5B | C3H10T1/2 | Shh HDL vs Shh HDL + Scube2 (50%) | 0,654±0,06 au vs 0,498±0,024 au, n=6 | <0,0001 |
| S5C | C3H10T1/2 | Shh HDL vs Shh HDL Scube2 | 6,114±1,085 vs 5,6±0,5225, n=5 | 0,3783 |

| | | | | |
|---|---|---|---|---|
| S5C | C3H10T1/2 | Shh HDL vs Shh HDL Scube2 | 2,088±0,46 vs 1,641±0,3147, n=5 | 0,1164 |
| S5D | C3H10T1/2 | Shh HDL vs 8908-SH HDL (100%) | 2,583±0,077 vs 1,263±0,079, n=6 | <0,0001 |
| S5D | C3H10T1/2 | Shh HDL vs $^{C25S}$ShhN HDL (100%) | 2,583±0,077 vs 0,3888±0,126, n=6 | <0,0001 |
| S5D | C3H10T1/2 | 8908-SH HDL vs $^{C25S}$ShhN HDL (100%) | 1,263±0,079 vs 0,3888±0,126, n=6 | <0,0001 |
| S5D | C3H10T1/2 | Shh HDL vs 8908-SH HDL (50%) | 1,468±0,1786 vs 0,6655±0,485, n=6 | <0,0001 |
| S5D | C3H10T1/2 | Shh HDL vs $^{C25S}$ShhN HDL (50%) | 1,468±0,1786 vs 0,319±0,075, n=6 | <0,0001 |
| S5D | C3H10T1/2 | 8908-SH HDL vs $^{C25S}$ShhN HDL (50%) | 0,6655±0,485 vs 0,319±0,075, n=6 | <0,0001 |
| S5D | C3H10T1/2 | mock | 0,416±0,054, n=6 | |
